# Facilitating Patient and Public Involvement in basic and preclinical health research

**DOI:** 10.1101/425371

**Authors:** James Maccarthy, Suzanne Guerin, Anthony G. Wilson, Emma R. Dorris

**Affiliations:** UCD Centre for Arthritis Research, School of Medicine, UCD Dublin, Ireland; UCD School of Psychology, UCD Dublin, Ireland

**Author notes:** **Corresponding Author:** Emma Dorris.

## Abstract

Involving patients in research broadens a researcher’s field of influence and may generate novel ideas. Preclinical research is integral to the progression of innovative healthcare. These are not patient-facing disciplines and implementing meaningful PPI can be a challenge. A discussion forum and thematic analysis identified key challenges of implementing PPI for preclinical researchers. In response we developed a “PPI Ready” planning canvas. For contemporaneous evaluation of PPI, a psychometric questionnaire and an open source tool for its evaluation were developed. The questionnaire measures information, procedural and quality assessment. Combined with the open source evaluation tool, researchers are notified if PPI is unsatisfactory in any of these areas. The tool is easy to use and adapts a psychometric test into a format familiar to preclinical scientists. Designed to be used iteratively across a research project, it provides a simple reporting grade to document satisfaction trend over the research lifecycle.

## Introduction

Involving patients or the interested public in research broadens a researcher’s field of influence, generating novel ideas, challenges and discussions. Basic, translational and preclinical research (hereto preclinical) is integral to the progression of innovative healthcare, indeed the majority of National Institute for Health (NIH) funds in the USA are focused on preclinical research (1). Preclinical research is not a traditionally patient-facing discipline and implementing meaningful patient and public involvement (PPI) can be a serious challenge in the absence of well-defined support structures.

Increasingly in healthcare, patients and the interested public are sought as partners in study design and governance. This trend is growing due to an increasing requirement by national, international and charitable funding bodies to include PPI as a condition of funding (2-6). We use the INVOLVE definition of PPI, whereby PPI in research is defined as research carried out with or by patients and those who have experience of a condition, rather than for, to, or about them (7). PPI has multiple demonstrated positive impacts on research including the potential to reduce waste in the research landscape (4, 8, 9). Furthermore, if a project or researcher is funded by a public body there is a duty to demonstrate accountability for the expenditure of supporting funding. Thus, incorporating PPI should be beneficial to both researcher and research outputs. There tends to be a recognition within the biomedical community that PPI can be beneficial to all stakeholders (10). Awareness of PPI is certainly increasing, however, the incorporation of valuable and meaningful PPI as standard research practice is progressing slower than expected(11, 12).

PPI is collaborative by its very nature and its impact on of its key stakeholders should be assessed in considering PPI success. Preclinical researchers are not patient-facing by nature and as such, some of the basic aspects of implementing meaningful PPI can be challenging without the relevant support structures. Understanding the perceived challenges and barriers as viewed by preclinical researchers is a first step in the development of appropriate resources and guidance documents to facilitate valuable collaboration and involvement between the public and the largest recipient cohort of publicly funded health research (13). Here we describe an analysis of the views of preclinical researchers into the challenges regarding PPI. In response, we have developed open source tools to help individual researchers prepare themselves to successfully incorporate PPI into their research, and have also developed a PPI assessment survey and analysis tool for iterative and contemporaneous assessment which facilitates refinement and improvement of PPI activities in response to feedback from the patients or public involved.

## Results and Discussion

### Determining preclinical researcher views on PPI

Basic, biomedical, biomolecular, translational and preclinical researchers were invited to attend a discussion forum focusing on what they perceive to be barriers or challenges to patient involvement in research. Researchers from all experience levels (undergraduate to PI), a range of disease areas and research disciplines were in attendance (researcher characteristics, table 1). The discussion forum started with an introduction to PPI, what it is, what it is not, how it applies to research, and consequences for funding, ethics and policy. A well-documented challenge for PPI is lack of standardised terms (14). This introduction set the terms for the discussion forum. The clarification of key terms was also printed and posted on the walls for the entirety of the discussion forum for reference to all who attended.

**Table 1:** PPI Discussion Forum Researcher Characteristics.

| Research Experience | n (%) | Primary Disease Area | n (%) | Primary Research Area | n (%) |
| --- | --- | --- | --- | --- | --- |
| Undergrad | 1 [6%] | Rheumatic Diseases | 4 [25%] | Statistical | 2 [13%] |
| Research Assistant | 1 [6%] | Diabetes | 2 [13%] | Molecular | 7 [44%] |
| Project Manager | 1 [6%] | Prostate Cancer | 4 [25%] | Basic | 6 [38%] |
| Postdoc | 7 [44%] | Breast Cancer | 2 [13%] | Psycho-social | 1 [6%] |
| Research Fellow | 3 [19%] | Cancer (Other) | 2 [13%] |  |  |
| Principal Investigator | 3 [19%] | Multi | 2 [13%] |  |  |

After the introduction the group broke into three groups. Each group had a facilitator and a note-taker. Attendees were also encouraged to note their opinions and views if they did not wish to vocalise them or if a point occurred to them after the discussion had moved on. Discussions focused on key challenges and facilitators for implementation of PPI. The challenges were broken into three concepts: barriers, worries and concerns. Barriers refer to institutional barriers that prevent or discourage active PPI (for example: ethics, funding, administrative supports). Worries refer to personal challenges a researcher faces in implementing PPI (for example: social anxiety, imposter syndrome). Concerns refer to challenges about the impact on research (for example: loss of research focus, public discomfort with research methods (such as animal research)).

Notes were collated and a textual and thematic analysis was performed (figure 1A). The major barriers identified were ethical challenges, engagement of patients and the interested public, funding to carry out PPI and a perceived lack of relevant guiding documents. Key concerns identified were time commitment, dilution of research objectives, sacrificing long-term value for short-term gains, and public judgement of ethically approved research (particularly animal models and experimentation). Personal worries that hindered PPI included lack of communication skills, fear of public disengagement, burden of expectation and social anxieties.

**Figure 1:**
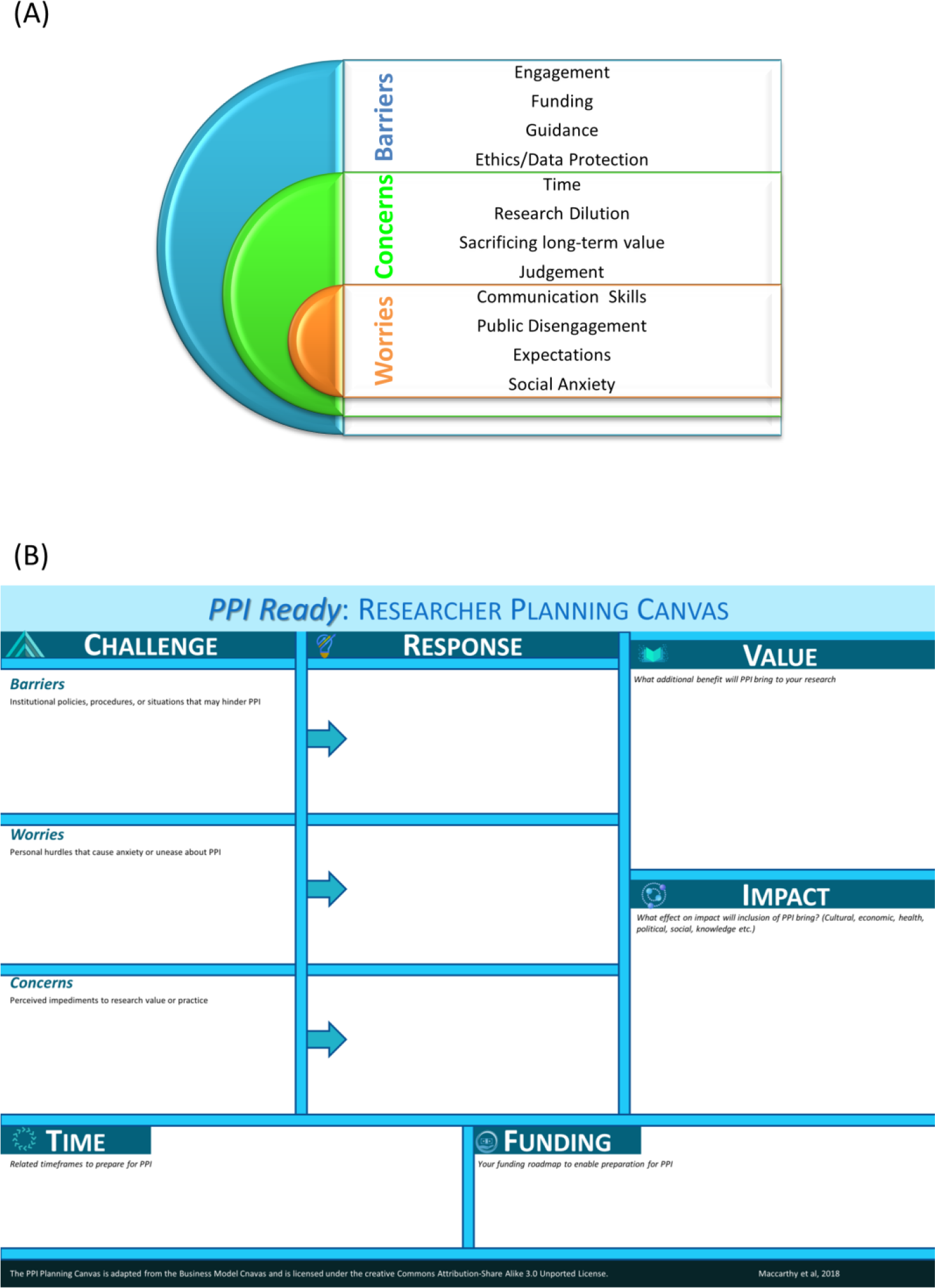
Key challenges hindering PPI and a tool for researchers to develop strategies to overcome these challenges. (A) Thematic analysis of a researcher discussion forum with preclinical researchers identified key challenges in the implementation and adoption of PPI into research practice. (B) A visual design tool based on the business model canvas, which we’ve called PPI Ready: Researcher Planning Canvas was developed to aid the individual researcher to explore their personal barriers to adopting PPI and map a strategy to overcome or circumvent them.

#### Communication is a key factor in researcher hesitancy to adopt PPI

A common theme across all challenges was communication. Communication barriers included a lack of predefined routes or mechanisms to source public or patient insight partners, and a lack of guidance documents regarding the legal and ethical responsibilities for local, national and international involvement of patients in research. “One of the biggest barriers is engaging patients and getting them to come in […] we need to be able to sell PPI to the public.”

Communication worries included discomfort or reluctance to talk about personal experiences with patients; fear of upsetting a patient; anxiety about how to handle disagreements with a patient insight partner: “Worried that a mistake I make will upset a patient”. Communication concerns included being able to explain and justify the need for preclinical and basic research, which has a longer term and less direct research impact. There was concern around public representatives bringing personal judgement upon research methods: ““Patients maybe against some aspects of research such as animal testing. We need […] to convey the importance of animal testing when it comes to the progression of research.” There was concern that the time spent justifying research that has already been through rigorous ethics application to public partners with alternative agendas could slow the research process and be counter-productive.

#### Communicating with vulnerable groups is not in the preclinical researchers’ toolkit

Preclinical research training may include scientific communications, presentation skills and even media training. Seldom, however, is training provided for communicating with vulnerable groups such as patient cohorts. The worry of appropriately communicating with vulnerable groups was particularly prominent with less experienced researchers and in researchers investigating fatal and life-limiting illnesses “The unfortunate reality of PPI is that some patients may have life-threatening diseases. What happens if a patient dies? As a researcher I don’t know what to do in that situation and I don’t know how to handle it”. More generally, there was a recurrent worry of upsetting the public, fear of “saying the wrong thing” or causing upset or public disengagement. The perceived lack of guidance, combined with the bespoke nature of PPI resulted in many researchers feeling the burden of publically representing their research and their institute. Whereas they may be comfortable presenting a fully peer-reviewed “sanctioned” piece of research output, there was a personal and professional vulnerability associated with sharing early stage and on-going research with the public. Researchers felt they would be guarded and could not discuss preliminary research fully if a public member was present for fear of said research being misconstrued or offering false hope.

#### Time consumption is a major concern hindering PPI implementation

The overwhelming largest concern about implementing PPI was the time required for the engagement, recruitment, procedural, and on-going management of public involvement. This may be a function of the relatively new strategy change towards PPI in Ireland. “I simply don’t have enough time to implement PPI. I can’t delegate any work in relation to PPI because I don’t think enough people know what it is and what to do.” In particular, researchers who had no experience in PPI had the strongest sense that the time consuming nature of PPI outweighed it’s benefit to preclinical research “Time spent engaging with patients is time spent away from doing research”. Again, this could be influenced by the lack of experience in engaging patients and the perceived lack of guidance and clarity around PPI (15) and dissociation between preclinical research and policy setting (16-18).

### Development of a tool to facilitate an individual researcher to be become PPI ready

PPI is bespoke; there is no standard formula for a good PPI initiative. However, one can be as prepared as possible to implement a bespoke PPI initiative. By reflecting on the main theoretical challenges for implementing PPI in advance of starting a research project, an individual researcher can facilitate the downstream success of their PPI initiative. An individual researcher’s theoretical challenges will stem from the uncertain boundaries of the concept of PPI. Therefore, we have adapted a standard business tool, the business model canvas (BMC), which is designed to enhance strategic thinking for business innovation and inform the exploitation of emerging opportunities in changing environments (19, 20). The PPI Ready Researcher Planning Canvas (figure 1B) is designed to enhance strategic thinking about PPI and enable a researcher to prepare themselves for PPI. The PPI Ready Canvas is a visual plan with elements describing a researchers challenges, proposed solutions, the solutions’ value proposition, research impact, financial and time costs. It is designed to assist researchers in aligning their activities by illustrating potential benefits and trade-offs.

The PPI Ready Researcher Planning Canvas elements allow researchers to consider what the challenges are that are hindering their implementation of PPI. It is broken down into Barriers: Institutional policies, procedures or situations that may hinder PPI; Worries: Personal hurdles that cause anxiety or unease about PPI; Concerns: Perceived impediments to research value or practice. There is corresponding space to consider the response to the challenges. In order to consider the cost/benefit analysis of implementing PPI, there is a module of Value: what additional benefit will PPI bring to your research; and Impact: What effect on impact will inclusion of PPI bring? Consider cultural, economic, health, political, social and knowledge impacts. To ground the proposed solutions or responses, there are modules to consider both the timeframe required to prepare for PPI and also the associated costs and funding required.

### A mechanism for iterative PPI assessment to allow incremental revisions and PPI development

Formalised PPI is a new concept in preclinical research. All initiatives, especially paradigm shifts, have associated risk. The earlier in the project lifecycle you can avoid a risk, the more accurate you can make your design. Many risks are not discovered until they have been to be integrated into a system and it is not feasible to identify all risks. Therefore, just like any area of the research cycle and design process, PPI should be assessed, evaluated and improved iteratively in response to experience and lessons learned. This may be tricky considering PPI by its very nature is not prescriptive. The form that PPI takes will be bespoke depending upon the needs of the research. However, the underlying concepts for valuable and meaningful PPI remain constant and can be measured and assessed.

We have developed a PPI satisfaction assessment survey to be used for iterative and contemporaneous PPI assessment throughout the life cycle of a research project (table 2). The survey should be completed by public/patient insight partners (PIPs) throughout their role in the lifecycle of the research project. This iterative approach has a number of benefits including: mitigated risks earlier because PPI elements can be integrated progressively; accommodation of changing requirements and tactics as experience is gained and lessons learned; facilitated refinement and improvement of PPI resulting in a more robust and meaningful initiative; facilitated knowledge exchange as institutions can learn from this approach and improve their process; and enhance reusability of PPI initiatives.

**Table 2:** Validated PPI Assessment Survey (PAS)

| In response to the following questions, please indicate your level of satisfaction. |  |  |
| --- | --- | --- |
| 0 = very dissatisfied, 5 = neither satisfied nor dissatisfied, 10 = very satisfied. |  |  |
| Value |  |  |
| Cate- |  |  |
| gory | Q. | How satisfied are you: |
| I | 1 | With the facilities provided? |
| I | 2 | That your needs were considered in the planning of the project meeting(s)? |
| P | 3 | With your level of contribution in relation to what you expected? |
| P | 4 | With your comfort in voicing your thoughts and opinions? |
| Q | 5 | With your understanding of the research project and its aims? |
| Q | 6 | That you understand the roles of other team members? |
| Q | 7 | With the communication and feedback tools that you have been asked to use in this project? |
| Q | 8 | That you know who to contact if an issue arises or, if you have concerns about the project or your involvement? |
| GAS | 9 | Overall, how satisfied/dissatisfied are you with your involvement in this project? |
| I= Informational; P=Procedural; Q=Quality; GAS= Global Assessment of Satisfaction |  |  |

#### PPI Assessment Survey Development

A review of the literature and qualitative review by patient insight partners (PIPs, n=12, table 3) lead to the development of 15 PPI assessment questions. Based on feedback from our PIPs and previous evaluation of scale types for health related research, we used an 11-point satisfaction scale with 3 anchor points (0= very dissatisfied, 5=neither dissatisfied nor satisfied, 10= very satisfied) (21, 22). The global assessment of satisfaction (GAS) question was added to the questionnaire as a measure of convergent validity. The 16 question PPI evaluation (PI) questionnaire and the well characterized 10-question generalized self-efficacy scale (GSE)(23) were piloted on a cohort of 62 patients or interested public who self-reported as having attended a meeting or event that gave them the opportunity to discuss or express their views about health research or to share their experience with researchers. 60/62 provided the appropriate consent for analysis. The analysis pathway is outlined in figure 2. Dimension reduction by oblique Factor analysis indicated multi-colinearity within the PI questionnaire (determinant 8.279×10^-11^). Sampling was adequate (Kaiser-Meyer-Olkin (KMO, 0.885)) and there was homogeneity of variance (Barlett’s Test df 105, p<0.001). Analysis of the correlation matrix (supplementary information) leads to the removal of highly collinear questions (Q3, 4, 7, 9, 11, 12 and 14). Factor analysis was rerun on the refined 8-question PI questionnaire (8QPI). Determinant value of 0.001 indicates an absence of collinearity. Sampling was adequate (KMO 0.886) and there is homogeneity of variance (Bartlett’s df 28, p<0.001).

**Table 3:**
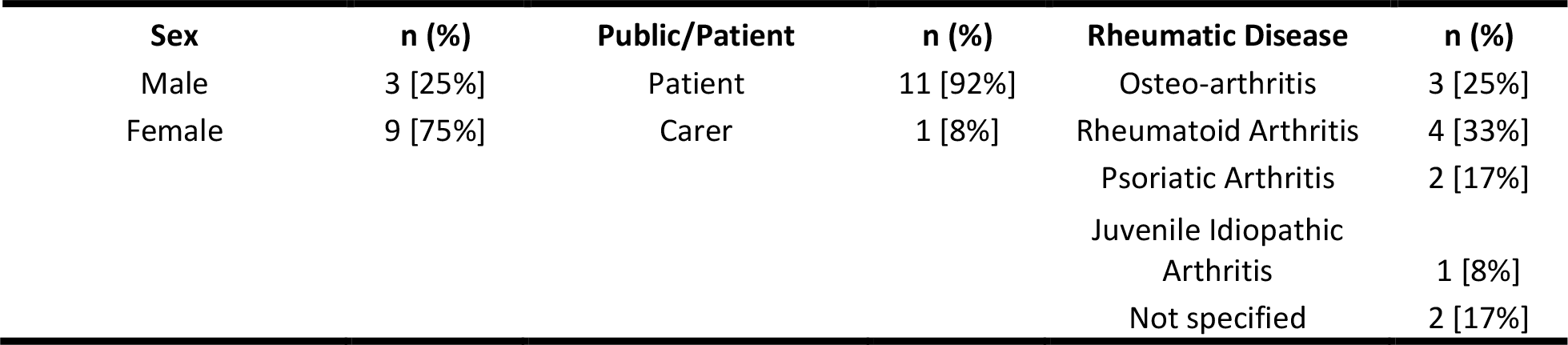
PIP Characteristics.

| Sex | n (%) | Public/Patient | n (%) | Rheumatic Disease | n (%) |
| --- | --- | --- | --- | --- | --- |
| Male | 3 [25%] | Patient | 11 [92%] | Osteo-arthritis | 3 [25%] |
| Female | 9 [75%] | Carer | 1 [8%] | Rheumatoid Arthritis | 4 [33%] |
|  |  |  |  | Psoriatic Arthritis | 2 [17%] |
|  |  |  |  | Juvenile Idiopathic Arthritis | 1 [8%] |
|  |  |  |  | Not specified | 2 [17%] |

**Figure 2:**
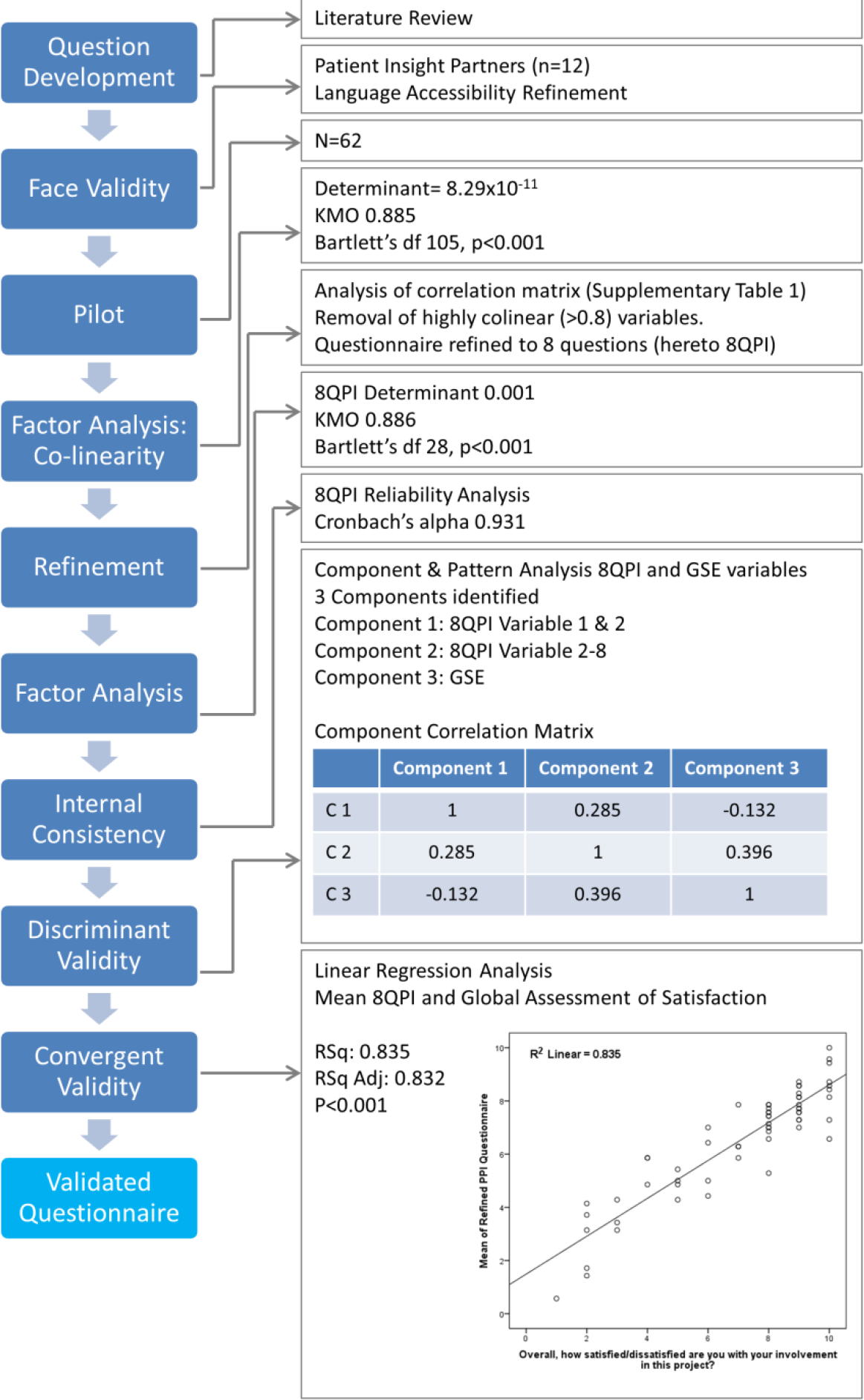
Development of the PPI assessment survey (PAS) Questions were adapted from the literature and reviewed by a patient insight partner (PIP) cohort of n=12 for face validity, relevance, jargon use, comprehension and language and format accessibility. Question wording and scale were refined in response and reassessed by a subset of the PIP cohort. The refined questionnaire and the general self-efficacy scale (GSE) were piloted on a cohort of patients and public that had attended a research involvement or engagement event. N=60 responses passed the consent check. Oblique factor analysis identified co-linearity between PPI questions which lead to question exclusion based on quantitative analysis and informed by qualitative input for PIPs. The refined questionnaire passed content validity analysis, including, internal validity, discriminant and convergent validity analysis.

Internal consistency was measured for 8QPI questionnaire via reliability analysis and produced a Cronbach’s alpha of 0.932, indicating high reliability/internal consistency of the variables in the scale. Discriminant validity was measured between the 8QPI and GSE scales. Factor analysis identified 3 components: GSE variables, 8QPI variable 1 and 2, and 8QPI variable 3-8 (supplementary information). Correlation between components ranged from −0.132-0.396 (figure 2). The variance extracted (range 0.56-0.68) is greater than the correlation squared (range 0.02-0.12) and thus discriminant validity is established.

Convergent validity was measured via the correlation between the per-case mean 8QPI response and the GAS question (Overall, how satisfied/dissatisfied are you with your involvement in this project?), which was included in the pilot survey. Linear regression demonstrated a positive correlation (R 0.914, RSq 0.835 and Adjusted RSq 0.832, figure 2).

#### An open-source simple analysis and reporting tool for the PPI assessment survey

Time consumption was highlighted as a major barrier to implementing PPI in the preclinical sciences. Therefore, development of a PPI assessment survey (PAS) would be redundant unless it was quick and easy to use. The tool is simple by design and at the suggestion of preclinical researchers who did not want to require any further training in order to be able to use it. An excel spreadsheet was selected as this programme is widely used and available and researchers were comfortable with it. We have therefore developed a series of simple excel templates with pre-formulated inputs for the analysis of the survey (figure 3). Furthermore, there are linked templates that can be sent to the PIP which allows the researcher’s file to be updated and analysed as the PIP fills in their survey response in a separate excel document.

**Figure 3:**
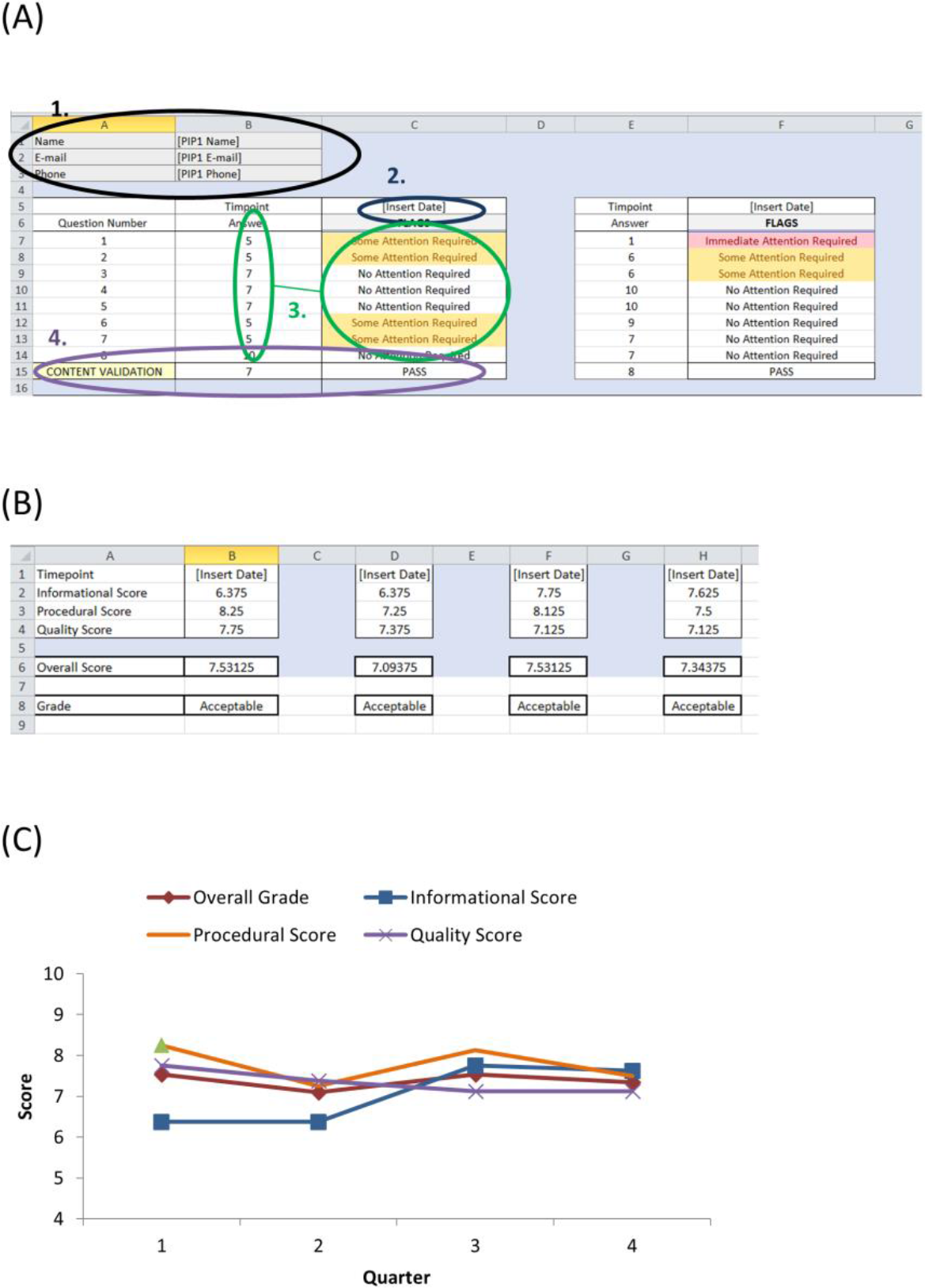
PAS assessment tool A preformatted excel file with embedded analysis formula is available for use with the PAS. (A) Tab 1 of the excel file is the data input tab. The PIP details are entered in (1). The date is entered in (2). (3) The responses to the PAS survey are entered in (3) and the flagging system is automatically generated. The PAS scores can be entered manually or can be updated via a linked excel file sent directly to the PIP(s). (4) The GSE score should be within two standard errors of the mean PPI scores (Q1-8). If not, a FAIL flag is generated and caution is advised in the interpretation of results. (B) Tab 2 is the output summary. It summarises the scores for each value category across all PIPs for each given time point and also generates an overall score and grade. (C) The scores can be plotted over time to visualise the PPI satisfaction trend. As illustrated, the overall score is relatively flat, but plotting of each value score illustrates that the informational assessment required attention and improved upon implementation of changes (after the second quarter) to increase clarity in communications.

The analysis is multi-dimensional and uses a flagging system, which preclinical researchers would be familiar with from standard quality control measures used by laboratory equipment. The analysis tool uses a flagging system for each question answered on an individual PIP level. The banding for our flag system was based upon a previously published large analysis of different response scales (24). If a PIP scored a question ≥7 no flag is generated. If a question is scored between 4-6 a yellow warning flag “some attention required” is generated. A score on any individual question of ≤3 generates a red warning flag “Immediate attention required”. These flags alert the researcher to risks associated with their PPI initiatives and allows them to put control measures in place. The iterative use of the PAS over time will facilitate trend analysis to determine the success of the control measures.

The GAS question Is included in the PAS (question 9). For validation of the survey during use, the analysis template has an inbuilt content validation (figure 3). If the GAS response is within the range of the Mean Q1-8 +/- two standard errors, the PAS is consistent in its measurement of satisfaction and a “PASS” flag is produced. If the GAS is outside this range a “FAIL” flag is produce and caution should be used in interpreting the results as there may be an underlying construct skewing the PAS response.

There is also an overall PAS assessment grade produced based on the mean PIP response across all questions. This can be used as an easy reporting measure for annual grant reports and for assessing a PPI initiative across the lifecycle of the project. The overall grade uses categorisation based upon standard risk assessment, a concept familiar to most preclinical researchers(25). An overall grade ≥7 is acceptable, 4-6 highlights a moderate risk to PPI success, and a grade <4 highlights a substantial risk to PPI success. The overall global grade is a useful tool for the measurement and reporting of PAS. However, if used in isolation there is a risk of masking potential subscale concerns. The PAS assesses three value levels: informational (quality and clarity of the information and communication provided, n= 2 questions), procedural fairness (consideration of the PIP needs, n=2 questions), and quality (how valued and valuable a PIP perceives the PPI initiative, n=4 questions). The analysis tool generates a per-category summary, which is the mean response from PIPs within each value category. This allows easy identification of improvements required between value categories and allows the researcher to pinpoint the area requiring refinement in order to improve their PAS grade (figure 3B).

#### Limitations of the PAS

The PAS is designed to facilitate assessment of PPI for preclinical researchers who tend to have little to no experience in qualitative, semi-quantitative or psychometric research. It is designed specifically for those with limited experience or access to established PPI resources. It addresses the key concepts underlying successful PPI collaborations between patients/public and researchers and ensures they are being addressed and assessed throughout the lifecycle of the project. The use of the open source analysis tool takes away the need for specialist training to use the PAS. The PAS can tell a researcher if there is a problem with PPI and in which category (information/communication, procedural fairness or quality); however, it does not identify the issue. The use of the flagging system on an individual PIP basis, allows for the rapid identification of whether it is an issue perceived by a single member of a PIP cohort, or if multiple PIPs have identified the same issue. This allows for simple direct follow-up with the PIP(s) to establish the cause of the issue without in depth qualitative analysis.

The circumvention of the qualitative analysis of other assessment tools, such as feedback forms which allow free text input, reduces the length of time and requirement for specialist knowledge and training for PPI assessment. It also allows the satisfaction trend to be easily tracked across time and in response to improvement measures. This can be beneficial but is also a limitation as there is a loss to the extent of information that can be assessed. A quantitative method cannot provide in-depth information on the motivations behind the levels of satisfaction nor provide details of PIP experiences with PIP (26). The use of the PAS with informal PIP feedback in response to issues identified in PAS analysis provides a realist and feasible solution given the researcher challenges described above. However, for the most robust development and progression of PPI as a research and impact tool, a mixed methods approach using the PAS and qualitative in-depth exploratory PPI narrative research and analysis could enable new topics and insights to emerge, which could also be confirmatory for the quantitative findings.

### Concluding Remarks

Preclinical researchers are increasingly required to wear multiple hats as part of the standard expectations of a research career position. Established roles include study design, conduct and experimental analysis but they are also now expected to be professional teachers, presenters, public speakers, writers, reviewers, editors, managers and mentors. There is increasing pressure to commercialise research in-house, act as innovators and become social and political influencers (27, 28). It is therefore unsurprising that time commitment, communication and lack of guidance were considered major challenges to the implementation of PPI in preclinical research. Understanding that researchers face personal challenges, anxieties and worries, as well as institution barriers and research concerns is a first step in encouraging meaningful PPI collaborations. Addressing the challenges faced by preclinical researchers will greatly help the development of resources and guidance documents to facilitate PPI to one of the largest recipient cohorts of publicly funded health research (1, 29).

The research describe here will aid preclinical researchers in the formulation of PPI. Adequate preparation in advance of a research project with public involvement should improve the success of that involvement. All new concepts have challenges and risk and involving the public in research is no different. Challenges are inevitable, however assessing public involvement iteratively through the use of the PAS described here within allows for those challenges to be addressed and managed contemporaneously. In conclusion, we have developed a tool specifically for the needs of the preclinical researchers to facilitate the implementation, assessment and improvement of PPI throughout the lifecycle of a research project.

## Methods

### Ethics Statement

Ethical approval from the local Institutional Review Board (UCD Human Research Ethics Committee). Survey responses were collected anonymously with informed consent of the participant for non-commercial use of the data provided.

### Patient and Public Involvement Statement

Patients were engaged through The Patient Voice in Arthritis Research PPI group. People with experience of living with any rheumatic disease from any area of Ireland were invited to apply to be involved in this study. Communication was remote, via email and phone. The patient insight partner group of 12 were sent a review guide and a structured template for their review (supplemental information). PIPs were asked specifically about: language accessibility, relevance, usefulness, necessity of the questionnaire, missing aspects, the scale used, and the likelihood of intended use, questionnaire length, overall views and alternative assessment methods. The questionnaire was adjusted in response to PIP feedback. Telephone follow-up was used to obtain views on questionnaire refinement. PIPs were also asked to share the survey pilot within their relevant networks. The results of the study will be shared with PIPs via email and through the patient/researcher co-produced newsletter of the UCD Centre for Arthritis Research, News Rheum.

### Researcher View on Public and Patient Involvement

Basic, translational and preclinical health researchers within the UCD College of Health and Agricultural Sciences were invited to a discussion forum to express their views on implementing PPI. All attendees consented to the non-commercial use of the data provided. A facilitated semi-structured discussion forum structure was used, consisting of three groups, each with a note-taker and a facilitator. There was also a roaming central facilitator. An option for written contribution was also provided for those who did not wish to voice their opinions. Notes were combined and a qualitative textual analysis was performed.

### Planning Canvas Development

In response to the preclinical researcher views on implementing PPI, we proposed that reflecting on the main theoretical challenges for implementing PPI, which stem from the uncertain boundaries of the concept, in advance of starting a research project would facilitate downstream success for PPI. A common tool in to help business rethink their business strategy in a fast-evolving landscape is the Business Model Canvas (BMC). The BMC is used to enhance strategic thinking about business innovation(20). We used the theory and design concept of the BMC informed by researcher views to develop the PPI Ready: Researcher Planning Canvas. The PPI Ready Canvas is designed to facilitate the researcher’s preparedness for PPI, rather than for planning an individual PPI activity.

### Questionnaire Development

A review of the health, public engagement, and marketing research literature was conducted (30-44). A long questionnaire of 15 questions was developed in response to the researcher discussion forum (supplemental information). Three key processes for assessment were information (n=4 questions), procedural fairness (n=4) and quality (n=7). The global assessment question “Overall, how satisfied/dissatisfied are you with your involvement in this project” was included as question 16 for convergent validity. Face Validity and Questionnaire Accessibility

A voluntary, community recruited panel of patient insight partners (PIP) (n=12) reviewed the questionnaire for face validity and language accessibility. The PIP review document can be found in supplementary methods. Questions were simplified and refined in response and changes discussed with PIPs. An 11 point satisfaction scale with 3 anchor points (at 0, 5 and 10) was used for all questions(22).

### Survey Pilot

A survey containing the 15 public involvement (PI) questions, one global assessment of satisfaction (GAS) question and the well characterized 10-question general self-efficacy (GSE) scale(23) was piloted on a cohort of 63 adults (full survey can be found in supplementary information). All respondents self-reported as having attended a meeting or event(s) that gave them the opportunity to discuss or express their views about health research or to share their experience with researchers. Three did not consent to data storage and use; therefore responses were excluded from analysis. Of the 60 respondents, 72% (n=43) were patients, 8% (n=5) were carers, 17% (n=10) were family members and 3% (n=2) were other members of the public. All surveys were complete and there were no missing entries.

### Data Analysis

Data was analysed in IBM SPSS v24. Factor analysis with correlation matrix was used to identify co-linearity and redundancy within the 15 PI questionnaire. As expected, there was a high degree of co-linearity and variables were removed based on a correlation greater than 0.8. To inform refinement, PIP insight, views, and reported relevance was considered. Eight questions remained after refinement (8QPI). Factor analysis was repeated on the 8QPI data, which met all requirements for determinant (>0.0001), KMO (>0.8) and Bartlett’s test for sampling adequacy (p<0.05). Cronbach’s alpha with a cut-off of 0.7 was used for internal consistency reliability testing. Discriminant validity was analysed via factor analysis and component analysis and correlation between 8QPI and GSE. Convergent validity was established via linear regression between the average 8QPI and GAS question, using a cut-off value of 1-(2SE) (two standard errors).

### Modelling the 8QPI into a Quality Control Framework

The validated PI questionnaire includes two questions for information assessment, two questions for assessment of procedural fairness, four questions for quality assessment and the GAS question. We developed a simple flagging system for each of the three key assessment categories that are measured on a per-individual basis. Furthermore, there is also an overall assessment grade that provides a global measure of PIP satisfaction with an individual PPI scheme.

The excel analysis template can be found in supplementary materials. PAS results are entered directly or via a linked file. For each question, responses of 0-3 issue an ‘Immediate Attention Required’ flag; 4-6 issued a ‘Some Attention Required’ flag and responses of 7-10 issued a ‘No Attention Required’ flag(24). Content validity is automatically tested by comparing the GAS response to the overall mean PAS (Q1-8) response. If the GAS is within the mean +/- 2 standard deviations (as determined in the PAS pilot), a ‘PASS’ flag is generated, otherwise a ‘FAIL’ flag is produced and the data must be interpreted with caution.

An output summary table is generated on tab 2. Each category (communication, procedural and quality) receives a score based on the mean responses for all PIPs. An overall score based on mean response for all questions (Q1-8) and associated PPI Grade is generated for simple PPI satisfaction reporting. Based on the risk matrix concept, overall scores of 0-3 are ‘High Risk’ for PIP dissatisfaction/PPI failure, scores of 4-6 represent a ‘Moderate Risk’ and a score of 7-10 are ‘Acceptable’(45).

## Acknowledgements

We wish to acknowledge our PIPs, all voluntary members of the UCD PPI initiative, The Patient Voice in Arthritis Research: Jacqui Browne, Wendy Costello, John Sherwin, Breda Fay, Eileen Tunney, Nicola Nestor and all Patient Voice in Arthritis Research contributors to the project.

The authors also wish to thank Dr. Geertje Schuitema (UCD College of Business) and Dr. Ricardo Segurado (UCD School of Public Health, Physiotherapy and Sports Science) for their assistance and advice.

## Funding Acknowledgement

Funding was provided by the Health Research Board of Ireland and the Irish Research Council.

## References

1. Moses H, Iii, Matheson DM, Cairns-Smith S, George BP, Palisch C, et al. The anatomy of medical research: Us and international comparisons. JAMA. 2015;313(2):174–89.

2. Ireland HRBo. Ireland: Research. Evidence. Action. HRB Strategy 2016–2020. Ireland: HRB; 2015.

3. van Thiel GS, Pieter. Priorities medicines for Europe and the world “A public health approach to innovation”. World Health Organization, 2013.

4. Richards T, Snow R, Schroter S. Co-creating health: more than a dream. BMJ. 2016;354.

5. Institute P-COR. What we mean by engagement USA: PCORI, 2015.

6. Research NIfH. Patients and the public. UK 2017.

7. INVOLVE. What is public involvement in research. UK: INVOLVE (NHS); [cited 2018 08 August 2018]; Available from: http://www.invo.org.uk/find-out-more/what-is-public-involvement-in-research-2/.

8. Chalmers I, Bracken MB, Djulbegovic B, Garattini S, Grant J, Gülmezoglu AM, et al. How to increase value and reduce waste when research priorities are set. The Lancet. 2014;383(9912):156–65.

9. Chu LF, Utengen A, Kadry B, Kucharski SE, Campos H, Crockett J, et al. “Nothing about us without us”—patient partnership in medical conferences. BMJ. 2016;354.

10. Pollock J, Raza K, Pratt AG, Hanson H, Siebert S, Filer A, et al. Patient and researcher perspectives on facilitating patient and public involvement in rheumatology research. Musculoskeletal Care. 2016;15(4):395–9.

11. Ocloo J, Matthews R. From tokenism to empowerment: progressing patient and public involvement in healthcare improvement. BMJ Quality & Safety. 2016.

12. Nagraj S, Gillam S. Patient participation groups. BMJ. 2011;342.

13. Chatterjee SK, Rohrbaugh ML. NIH inventions translate into drugs and biologics with high public health impact. Nature Biotechnology. 2014;32:52.

14. Liabo K, Boddy K, Burchmore H, Cockcroft E, Britten N. Clarifying the roles of patients in research. BMJ. 2018;361.

15. Patricia Wilson EM, Julia Keenan, Elaine McNeilly, Claire Goodman, Amanda Howe, Fiona Poland, Sophie Staniszewska, Sally Kendall, Diane Munday, Marion Cowe, and Stephen Peckham. ReseArch with Patient and Public invOlvement: a RealisT evaluation – the RAPPORT study. Health Services and Delivery Research, No 338. Southampton (UK): NIHR Journals Library; 2015.

16. de Jong SPL, Smit J, van Drooge L. Scientists’ response to societal impact policies: A policy paradox. Science and Public Policy. 2016;43(1):102–14.

17. Boswell C, Smith K. Rethinking policy ‘impact’: four models of research-policy relations. Palgrave Communications. 2017;3(1):44.

18. Gianos PL. Scientists as Policy Advisers: the Context of Influence. Western Political Quarterly. 1974;27(3):429–56.

19. Osterwalder Alexander PY. Business Model Generation: A Handbook for Visionaries, Game Changers, and Challengers: Wiley; 2010.

20. Spieth P, Schneckenberg D, Ricart JE. Business model innovation – state of the art and future challenges for the field. R&D Management. 2014;44(3):237–47.

21. Courser ML, Paul J. Item-Nonresponse and the 10-point response scale in telephone surveys. Survey Practice. 2012.

22. Preston CC, Colman AM. Optimal number of response categories in rating scales: reliability, validity, discriminating power, and respondent preferences. Acta Psychologica. 2000;104(1):1–15.

23. Jerusalem RSaM. Generalized Self-Efficacy Scale. Windsor, England: NFER-Nelson; 1995.

24. van Beuningen JvdH, Karolijne; Moone, Linda. Measuring well-being. An analysis of different response scales. The Hague, Netherlands: Statistics Netherlands; 2014.

25. Lyon BKP, Georgi The art of assessing risk. Professional Safety. 2016:40–51.

26. Chow MYK, Quine S, Li M. The benefits of using a mixed methods approach – quantitative with qualitative – to identify client satisfaction and unmet needs in an HIV healthcare centre. AIDS Care. 2010;22(4):491–8.

27. Arnette R. Wearing Many Hats. Science Magazine. 2005.

28. Hollenbach AD. The many hats of an academic researcher. ASBMB Today. 2014.

29. Moses H, Martin JB. Biomedical Research and Health Advances. New England Journal of Medicine. 2011;364(6):567–71.

30. Staniszewska S, Brett J, Mockford C, Barber R. The GRIPP checklist: Strengthening the quality of patient and public involvement reporting in research. International Journal of Technology Assessment in Health Care. 2011;27(4):391–9. Epub 2011/10/17.

31. Stocks SJ, Giles SJ, Cheraghi-Sohi S, Campbell SM. Application of a tool for the evaluation of public and patient involvement in research. BMJ Open. 2015;5(3).

32. Edelman N, Barron D. Evaluation of public involvement in research: time for a major re-think? Journal of Health Services Research & Policy. 2016;21(3):209–11.

33. Crocker JC, Boylan A-M, Bostock J, Locock L. Is it worth it? Patient and public views on the impact of their involvement in health research and its assessment: a UK-based qualitative interview study. Health Expectations. 2017;20(3):519–28.

34. Staley K. ‘Is it worth doing?’ Measuring the impact of patient and public involvement in research. Research Involvement and Engagement. 2015;1(1):6.

35. Choy E, Perrot S, Leon T, Kaplan J, Petersel D, Ginovker A, et al. A patient survey of the impact of fibromyalgia and the journey to diagnosis. BMC Health Services Research. 2010;10(1):102.

36. Lempp HK, Hatch SL, Carville SF, Choy EH. Patients’ experiences of living with and receiving treatment for fibromyalgia syndrome: a qualitative study. BMC Musculoskeletal Disorders. 2009;10:124-.

37. Raymond MC, Brown JB. Experience of fibromyalgia. Qualitative study. Canadian Family Physician. 2000;46:1100–6.

38. Rodham K, Rance N, Blake D. A qualitative exploration of carers’ and ‘patients’ experiences of fibromyalgia: one illness, different perspectives. Musculoskeletal Care. 2010;8(2):68–77.

39. Connolly M, McLean S, Guerin S, Walsh G, Barrett A, Ryan K. Development and Initial Psychometric Properties of a Questionnaire to Assess Competence in Palliative Care: Palliative Care Competence Framework Questionnaire. American Journal of Hospice and Palliative Medicine®. 2018:1049909118772565.

40. Kernohan WG, Brown MJ, Payne C, Guerin S. Barriers and facilitators to knowledge transfer and exchange in palliative care research. BMJ Evidence-Based Medicine. 2018;23(4):131.

41. Lin F-H, Tsai S-B, Lee Y-C, Hsiao C-F, Zhou J, Wang J, et al. Empirical research on Kano’s model and customer satisfaction. PloS one. 2017;12(9):e0183888.

42. Boynton PM, Greenhalgh T. Selecting, designing, and developing your questionnaire. BMJ: British Medical Journal. 2004;328(7451):1312–5.

43. Rowe G, Frewer LJ. A Typology of Public Engagement Mechanisms. Science, Technology, & Human Values. 2005;30(2):251–90.

44. Perlaviciute G, Schuitema G, Devine-Wright P, Ram B. At the Heart of a Sustainable Energy Transition: The Public Acceptability of Energy Projects. IEEE Power and Energy Magazine. 2018;16(1):49–55.

45. Baybutt P. Guidelines for designing risk matrices. Process Safety Progress. 2017;37(1):49–55.

